# Conserved population dynamics in the cerebro-cerebellar system between waking and sleep

**DOI:** 10.1101/2022.05.06.490891

**Authors:** Wei Xu, Felipe De Carvalho, Andrew Jackson

## Abstract

Despite the importance of the cerebellum for motor learning, and the recognised role of sleep in motor memory consolidation, surprisingly little is known about activity in the sleeping cerebro-cerebellar system. Here we used wireless recording from M1 and the cerebellum in monkeys to examine the relationship between patterns of single-unit spiking activity observed during waking behaviour and in natural sleep. Across the population of recorded units, we observed similarities in the timing of firing relative to local field potential features associated with both movements during waking and up-states during sleep. We also observed a consistent pattern of asymmetry in pair-wise cross-correlograms, indicative of preserved sequential firing in both wake and sleep at low frequencies. Despite the overall similarity in population dynamics between wake and sleep, there was a global change in the timing of cerebellar activity relative to motor cortex, from contemporaneous in the awake state, to motor cortex preceding the cerebellum in sleep. We speculate that similar population dynamics in waking and sleep may imply that cerebellar internal models are activated in both states, but that their output is decoupled from motor cortex in sleep. Nevertheless, spindle frequency coherence between the cerebellum and motor cortex may provide a mechanism for cerebellar computations to influence sleep-dependent learning processes in the cortex.

**Significance statement:** It is well known that sleep can lead to improved motor performance. One possibility is that synaptic changes during sleep result from off-line repetitions of neuronal activity patterns in brain areas responsible for the control of movement. In this study we show for the first time that neuronal patterns in the cerebro-cerebellar system are conserved during both movements and sleep up-states, albeit with a shift in the relative timing between areas. Additionally, we show the presence of simultaneous M1-cerebellar spike coherence at spindle frequencies associated with up-state replay and postulate that this is a mechanism whereby a cerebellar internal models can shape plasticity in neocortical circuits during sleep.

## Introduction

The motor system must learn to generate complex, skilled movements, but the old adage of ‘practice makes perfect’ has in recent decades been updated to ‘practice with sleep makes perfect’ (Walker, Brakefield et al. 2002, Walker and Stickgold 2005, Fogel and Smith 2006, Fogel, Smith et al. 2009, Menicucci, Piarulli et al. 2020). The cerebellum has long been implicated in the process of motor learning (De Zeeuw 2021) but surprisingly, the possible involvement of the cerebellum in sleep-dependent consolidation and overnight performance improvements has been largely ignored (Canto, Onuki et al. 2017). Although sleep is known to benefit procedural as well as episodic memories (Nishida and Walker 2007, Diekelmann and Born 2010, Fogel, Albouy et al. 2017), research into the mechanisms of sleep-dependent learning has generally focussed on oscillatory interactions between the hippocampus and neocortex. In particular, replay of waking patterns of population activity, associated with nested spindles and slow oscillations, is thought to constitute a mechanism by which short-term episodic memories in the hippocampus may be consolidated into long-term storage in the neocortex (Staresina, Bergmann et al. 2015, Ngo, Fell et al. 2020). However, sleep spindle density also correlates with consolidation of motor sequence learning, a task associated with a progressive shift from cerebellar to cortico-striatal activity across multiple practice sessions (Doyon, Song et al. 2002, Spampinato, Celnik et al. 2020). This raises the intriguing possibility that sleep, and perhaps sleep oscillations, may play a similar role in transferring procedural memory traces from the cerebellum to the neocortex.

We have recently shown that the cerebellum is an active participant in sleep and sleep oscillations (Xu, De Carvalho et al. 2021). Functional connectivity analysis applied to local field potentials revealed causality directed from motor cortex to the cerebellum at low frequencies. By contrast, at spindle frequencies during identified up-state events we observed causality directed from the cerebellum to motor cortex, via the thalamus. This finding offers a tantalising clue to a potential involvement of cerebro-cerebellar circuits in off-line learning processes. Low-frequency patterns of activity within motor cortex at the onset of up-states resemble those seen during waking movements (Hall, de Carvalho et al. 2014). Moreover, sleep spindles are associated with cortical up-states and periods of enhanced plasticity (Andrillon, Nir et al. 2011). Thus, it is possible that cerebellar output in sleep influences cortical plasticity via spindle-frequency communication through the thalamus.

To better understand oscillatory interactions between the cerebellum and neocortex during sleeping and waking, we compared activity patterns at the single cell and population level within and between functionally connected areas of primary motor cortex and the cerebellum. We used time-domain cross-correlation and frequency-domain cross-spectral measures to characterise low-frequency dynamics associated with movements and up-states. We show these dynamics are broadly conserved between waking and sleep, although in the awake state M1 and cerebellar spiking activity is, on average, synchronous whereas during sleep there is a relative time-lag from the cortex to cerebellum. We propose that this may reflect a changing role for the cerebellum, from an active participant in predictive control of waking movement to predicting the consequences of fictive movements in sleep, which could constitute a mechanism for off-line consolidation of skill learning.

## Materials and methods

### Experimental design and statistical analysis

Three female rhesus macaques (U: 7 years old, 7.7kg; T: 10 years old, 7.8kg; Y: 6 years old, 6.9kg; housed in pairs) were used in this study. Recordings from these monkeys have previously been reported in two other studies (Xu, de Carvalho et al. 2019, Xu, De Carvalho et al. 2021). Experimental objectives and procedures were approved by the local Animal Welfare Ethical Review Board and licensed by the UK Home Office in accordance with the Animals Scientific Procedures Act of 1986 (revised in 2013).

Surgeries were performed to implant a titanium head casing, fixed linear microelectrode arrays (16-channel LMA, 12.5 μm platinum-iridium, 500 kΩ, MicroProbes for Life Sciences, USA) and 12 moveable flexible microwires in the right M1 (hand area) and 8 in the left cerebellum (lobules IV/V intermediate zone). These two areas are known to be anatomically connected (Kelly and Strick 2003). Signals were referenced to a low impedance microwire placed over the M1 dura. As previously described (Xu, De Carvalho et al. 2021), cerebellar microwires were preloaded into a 16-gauge guide-needle in order to penetrate the tentorium cerebelli. Both local field potentials (LFPs) and action potentials are derived from recordings from the flexible microwires. Recordings from new neurons were obtained by moving the flexible microwires under light sedation with ketamine approximately once per week.

In monkey U we recorded electromyogram (EMG) from six arm muscles on the left (extensor carpi radialis, extensor carpi ulnaris, flexor carpi radialis, flexor carpi ulnaris, biceps, triceps) using pairs of insulated stainless-steel wires (AS632, Cooner wire, USA) sutured into the muscle and tunnelled subcutaneously to the head casing.

Our free-behaving dataset for monkeys U, T and Y comprised 103, 29 and 17 sessions respectively, containing 438, 26 and 52 M1 spikes and 96, 14 and 35 cerebellar spikes. A subset of these sessions had simultaneously recorded M1 and cerebellar spikes (U: 49, T: 7, Y: 16). The total number of M1-M1, Cb-Cb and M1-Cb spike pairings were 1292, 85, and 590 across three animals. 22 sessions from monkey U contained simultaneous M1 and EMG recordings. Cerebellar spikes in this data set have previously been shown to consist of overwhelmingly Purkinje cell simple spikes by virtue of their similarity with a subset of neurons with identifiable complex spikes (Xu, De Carvalho et al. 2021).

Additionally monkeys U and Y were both trained in the lab to move an on-screen cursor from centre of screen to one of eight targets on the periphery by generating isometric flexion-extension (vertical) and radial-ulnar (horizontal) torque with wrist restrained in a pronated position. This task has previously been described in detail (Hall, de Carvalho et al. 2014). Our dataset includes 56 task sessions for monkey U, yielding 329 M1 and 268 cerebellar neurons (1657 simultaneous pairs M1-Cb) and 8 sessions for monkey Y yielding 83 M1 and 10 cerebellar neurons (114 simultaneous M1-Cb pairs).

Pair-wise metrics in our study (e.g. relative phases and cross correlation asymmetries – see below) are not statistically independent (since an individual cell can contribute to multiple cell pairs), so the standard parametric approach to testing the significance of their correlation is invalid. Therefore, in order to test whether there was a significant correlation between relative phase and cross correlation asymmetries (across cell pairs) in waking and sleep, we used our previous non-parametric approach (Xu, de Carvalho et al. 2019). This involved generating 1000 surrogate datasets by randomly shuffling the labelling of cells within each session and across sessions with the same number of cells to bootstrap the distribution of correlation coefficients under the null hypothesis that there was no relationship between wake and sleep. This distribution was used to derive upper and lower significance thresholds to reject the null hypothesis at P<0.05 in a two-sided test.

### Data recording

Recordings were made using a battery-powered, wearable data-logger developed in-house and described previously (Xu, de Carvalho et al. 2019). The device was based on two RHD2132 Intan bio-amplifiers (Intan Technologies, USA) and a STM32 F407 microcontroller (STMicroelectronics, Switzerland) that streamed data onto a 32GB microSD card. Signals from microwires were sampled at 20kHz. Signals from EMGs were sampled at 1000Hz. For sessions containing simultaneous M1 and cerebellar microwire recordings we recorded from 5 M1 microwires and 3 cerebellar microwires. For sessions containing only M1 recordings we recorded from 8 M1 microwires. The device also recorded field potentials from rigid linear multi-electrode arrays (Microprobes for Life Science, USA) but this data does not feature in this study. Recording sessions lasted around 20 hours and included a full night’s sleep and waking free behaviour periods.

### Signal processing

All signal processing and data analysis were carried out using MATLAB (MathWorks USA). All off-line filtering used 4-pole Butterworth filters applied in forward and reverse directions. M1 local field potentials (LFPs) were derived from the mean of down-sampled (20kHz to 250Hz, after anti-aliasing filtering) signals recorded from microwires with a low impedance wire on the dura over M1 as a reference. The referenced signals from all M1 microwires were averaged to yield a single mean M1 LFP. Spikes were discriminated after high-pass filtering raw signals above 300Hz using principal component analysis and clustering. Bipolar EMG signals were high-pass filtered at 50Hz, rectified, down-sampled (1000Hz to 250Hz) and averaged across all six arm muscles. All frequency domain analysis used 1024 sample FFT windows.

### Removal of REM and arousal periods during sleep

Periods of putative REM and arousal were removed from sleep data using our previously published and validated method (Xu, De Carvalho et al. 2021): seep recordings were divided into 30s long windows and those containing high gamma power (50-125Hz) above 1.13 times the average over the entire sleep period were removed. Note this method cannot distinguish between REM and arousal periods and therefore both are excluded from the dataset. Therefore the term ‘sleep’ in this study refers only to nREM sleep.

### Identification of sleep up-states

To identify sleep up-states we employed a widely used method (Nir, Staba et al. 2011, Xu, De Carvalho et al. 2021) whereby the LFP was filtered between 0.5-4Hz and negative-going halfwaves with zero-crossings between 0.25-1s were identified. The top 20% of these half-waves in terms of magnitude were selected for further analysis.

### Cell-cell coherence and relative phase

Spike times were binned using 4ms wide bins to produce a signal with the same sampling rate as LFPs and EMGs (250Hz). Coherence between pairs of cells (M1-M1, Cb-Cb and M1-Cb), as well spike-EMG coherence was calculated using:

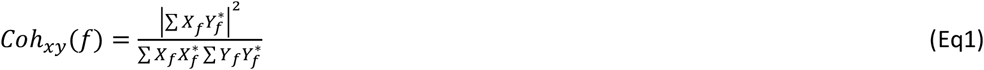

where *X*_*f*_ and *Y*_*f*_ are the Fourier coefficients of signals x and y, calculated over non-overlapping 1024 point windows with a frequency resolution of 0.24Hz. * denotes complex conjugate.

We used the phase of the cross-spectrum to quantify the phase relationship of activity at a given frequency between every cell pair. We were interested in whether these phase relationships were consistent across the population between waking and sleep. To do this we calculated the circular-circular correlation coefficient between relative phase in wake and sleep across all cell pairs at a given frequency:

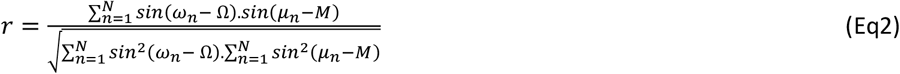

where r is the circular-circular correlation coefficient, N is the number of pairs, ω and µ denote the relative phase between each pair in waking and sleep respectively, and Ω and M denote the circular mean of these relative phases.

### Spike-LFP cross-correlation phase and cell-cell cross-correlation asymmetry

Time-domain cross-correlations between M1/Cb cell activity and M1 LFP were calculated using the firing rates binned at 250 Hz for relative lags up to ±2 s, low-pass filtered at 2 Hz. A Hilbert transform was used to calculate the phase of the cross-correlation at time zero. Consistency of spike-LFP relative phase across the population in wake and sleep was again assessed using circular-circular correlation.

We also calculated time-domain cross-correlations between the firing of M1-M1, Cb-Cb and M1-Cb cell pairs for relative lags up to ±1 s. Cross-correlation asymmetry was calculated from the difference in area under the cross-correlation under the left and right side of zero-lag, divided by the total area under the curve. Consistency of cell-cell cross-correlation asymmetry across the population in wake and sleep was assessed using linear correlation.

## Results

### Wireless recording during free movement and sleep

We used a wearable device to record single units in the primary motor cortex (M1), contralateral cerebellum (Cb) and contralateral forearm muscle EMGs. For some sessions we recorded neurons in M1 (161 neurons over 22 sessions) simultaneously with EMGs, and in another 72 sessions we recorded simultaneously from both M1 (289 neurons) and cerebellum (145 neurons). Examples of neuronal signals are shown in Figure 1A. Simultaneous recordings suggested that M1 firing rates were modulated with arm EMG throughout the recording (Figure 1B). We confirmed this by calculating coherence spectra between the firing of individual M1 neurons and rectified EMG (Figure 1C). This revealed a prominent peak at ∼1Hz reflecting the dominant spectral content of voluntary movement during waking, which was notably absent during sleep.

**Figure 1.**
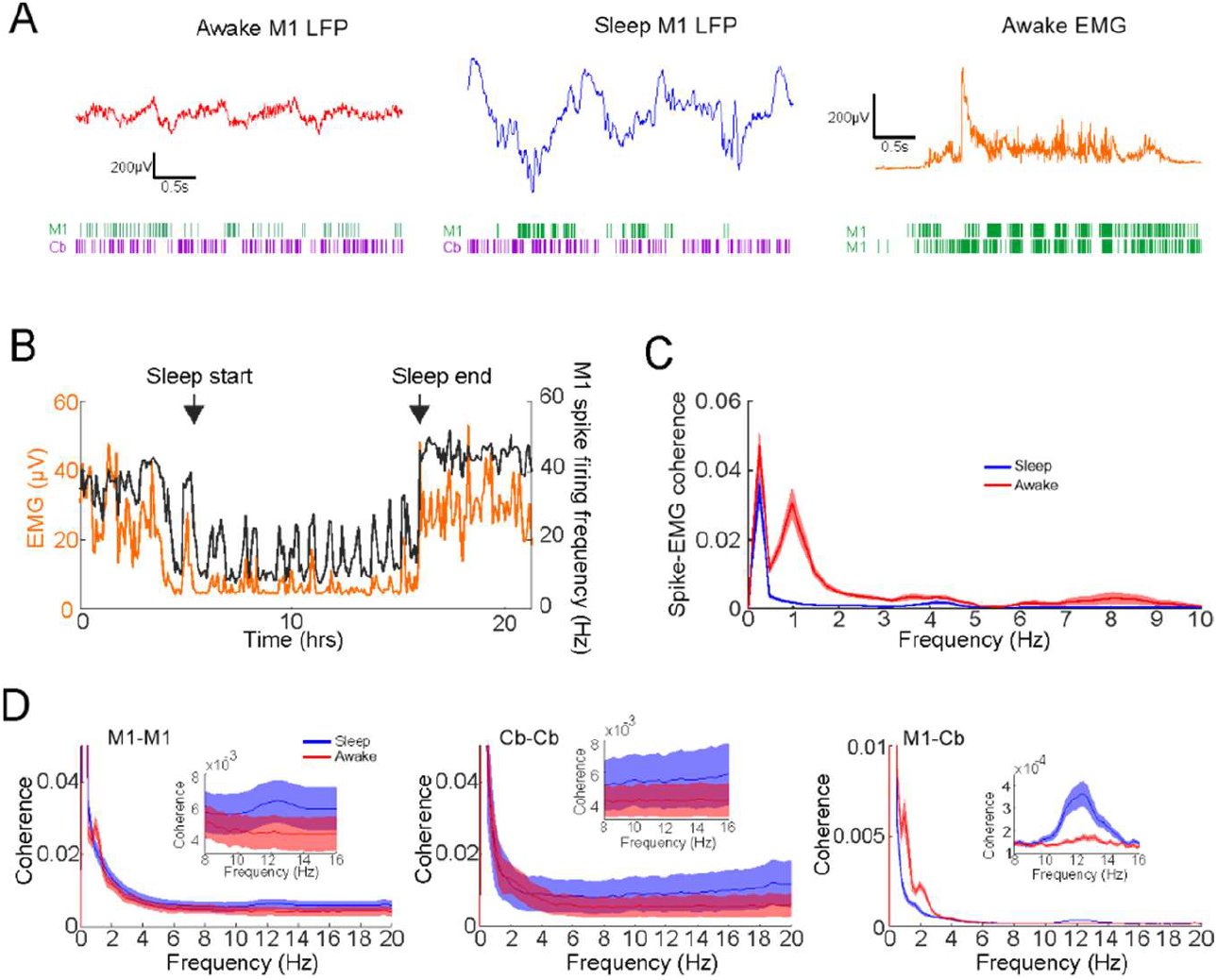
Coherence between firing rates and EMG during free behaviour and sleep. A. Example of M1 LFPs recorded simultaneously with M1 and cerebellar spiking (left and middle panel) and example of mean rectified EMGs from upper-limb muscles recorded simultaneously with M1 spiking. B. Example of mean firing rate and EMG recorded over an entire session. C. Mean coherence between M1 spiking and EMG during sleep and awake. D. Mean spike coherence between M1-M1, Cb-Cb and M1-Cb cell pairs. Inset shows coherence in the spindle frequency band.

Next, we calculated coherence between the firing rate of all neurons within the same area (M1-M1, Cb-Cb) and between areas (M1-Cb; Figure 1D). The inset panels show coherence in the spindle frequency band. Note that there was coherence within and between M1 and the cerebellum at low frequencies in both waking and sleep, although the inter-area M1-Cb low-frequency coherence was reduced in sleep. By contrast, coherence between M1 and Cb at spindle frequencies was enhanced in sleep. These firing rate results corroborate our previous analysis of LFPs in revealing oscillatory coupling between the cortex and cerebellum during sleep (Xu, De Carvalho et al. 2021).

### Awake movement events and sleep up-states are associated with high M1 firing rates and depth-negative LFP troughs

In order to examine in more detail neural activity associated with movement, we band-pass filtered the rectified EMG signals around the frequency of the coherence peak (0.5-2Hz; Fig. 1C) and selected the time of prominent peaks (2 SDs above mean peak amplitudes) as indicators of movement events to align M1 spike firing and LFP. M1 spike events aligned by movement events revealed a prominent firing rate peak that preceded the EMG peak (low pass filtered at 2Hz, Figure 2A) by an average of 92 ± 19 ms (± s.e.m), consistent with previous reports (Holdefer and Miller 2002). The peak in average M1 firing was also associated with a prominent trough in the M1 LFP (Figure 2B), consistent with previous studies (Destexhe, Contreras et al. 1999). Note however that the time at which individual neurons fired maximally relative to the EMG peak and LFP trough varied across neurons (Figure 2C). Such sequential firing of M1 neurons during movements has been reported previously (Churchland, Cunningham et al. 2012, Russo, Bittner et al. 2018, Xu, de Carvalho et al. 2019).

**Figure 2.**
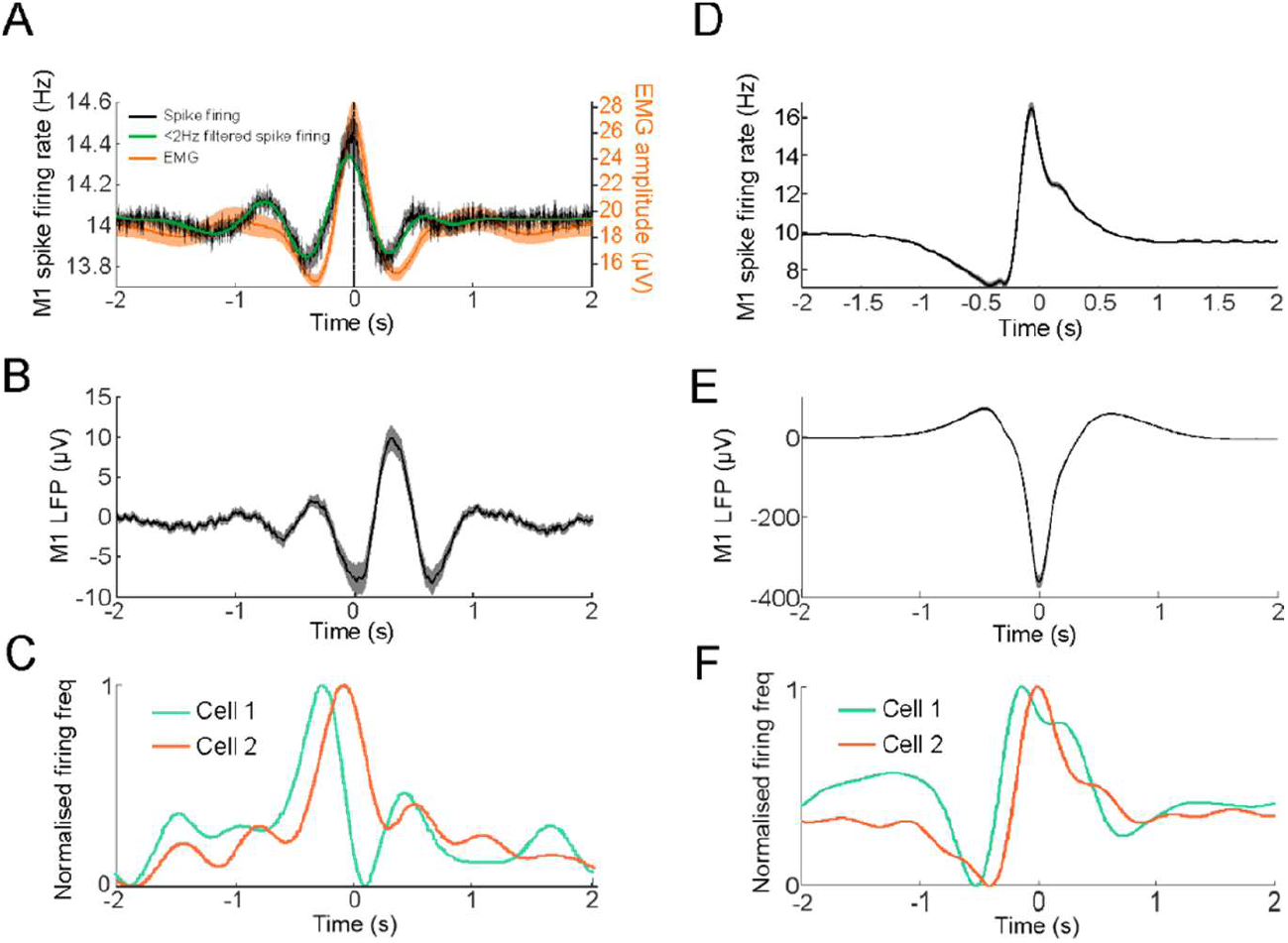
Sequential spiking activity during movements and sleep up-states. A. Waking EMG peak aligned mean M1 spike firing rate. B. M1 LFP aligned by EMG peaks. C. Example mean firing rates of two M1 neurons aligned by EMG peaks (low-pass filtered at 2Hz). D. Sleep up-state aligned mean M1 spike firing rate. Shaded region represent s.e.m. of differences in firing rates from background. E. Mean M1 LFP aligned by up-states. Shaded regions represent s.e.m. F. Example mean firing rates of two M1 neurons aligned by sleep up-states (low-pass filtered at 2Hz).

During non-REM sleep, LFP troughs and increased firing rates have been associated with up-states of the neocortical slow oscillation (Dickey, Sargsyan et al. 2021, Xu, De Carvalho et al. 2021). We used a well-established method to identify sleep up-states from M1 LFPs (Nir, Staba et al. 2011, Xu, De Carvalho et al. 2021) and confirmed that M1 spiking did indeed increase during sleep up-states in association with large negativities in the LFP (Figure 2D,E). As with activity during movement events, individual neurons showed firing rate profiles that peaked at different times relative to this LFP trough (Figure 2F).

### Similar low-frequency spike-LFP coupling between waking and sleep

We were interested to see if the general sequence of neuronal firing was preserved across the population between waking and sleep. We exploited the fact that neuronal activation in both waking and sleep was associated with an LFP trough, such that the shape of the spike-triggered LFP average gives an indication of whether individual neurons tend to fire earlier or later in relationship to that trough. M1 and cerebellar (Cb) spikes were both phase-locked to low frequency M1 LFPs – as shown by the large low-frequency component in spike-triggered averages of M1 LFPs (Figure 3A, B left). As expected, when averaged across all neurons, the time of spiking was associated with an LFP trough in both waking and nREM sleep. However, across the population there was considerable variability in this profile. Figure 3A (right) shows low-pass (<2 Hz) filtered spike-triggered averages of M1 LFP for all M1 and Cb cells. The cells are ordered according to the phase of the cell-LFP relationship, measured from the phase of the Hilbert transform at time-zero. Figure 3B (right) shows the awake spike-triggered averages plotted in the same order (i.e. derived from the sleep data). It is apparent through visual inspection that there is similarity between the patterns at the population level. Cells that fire in advance of the LFP trough in sleep similarly do so in the awake data and vice versa. We quantified this similarity by calculating the circular-circular correlation coefficient between the phases (at time 0) of the cell-LFP relationship in waking and sleep across cells. This correlation was significantly positive for both M1 and Cb neurons (Figure 3C,D) suggesting that there is similarity in the population activity relative to LFP features in both waking and sleep.

**Figure 3.**
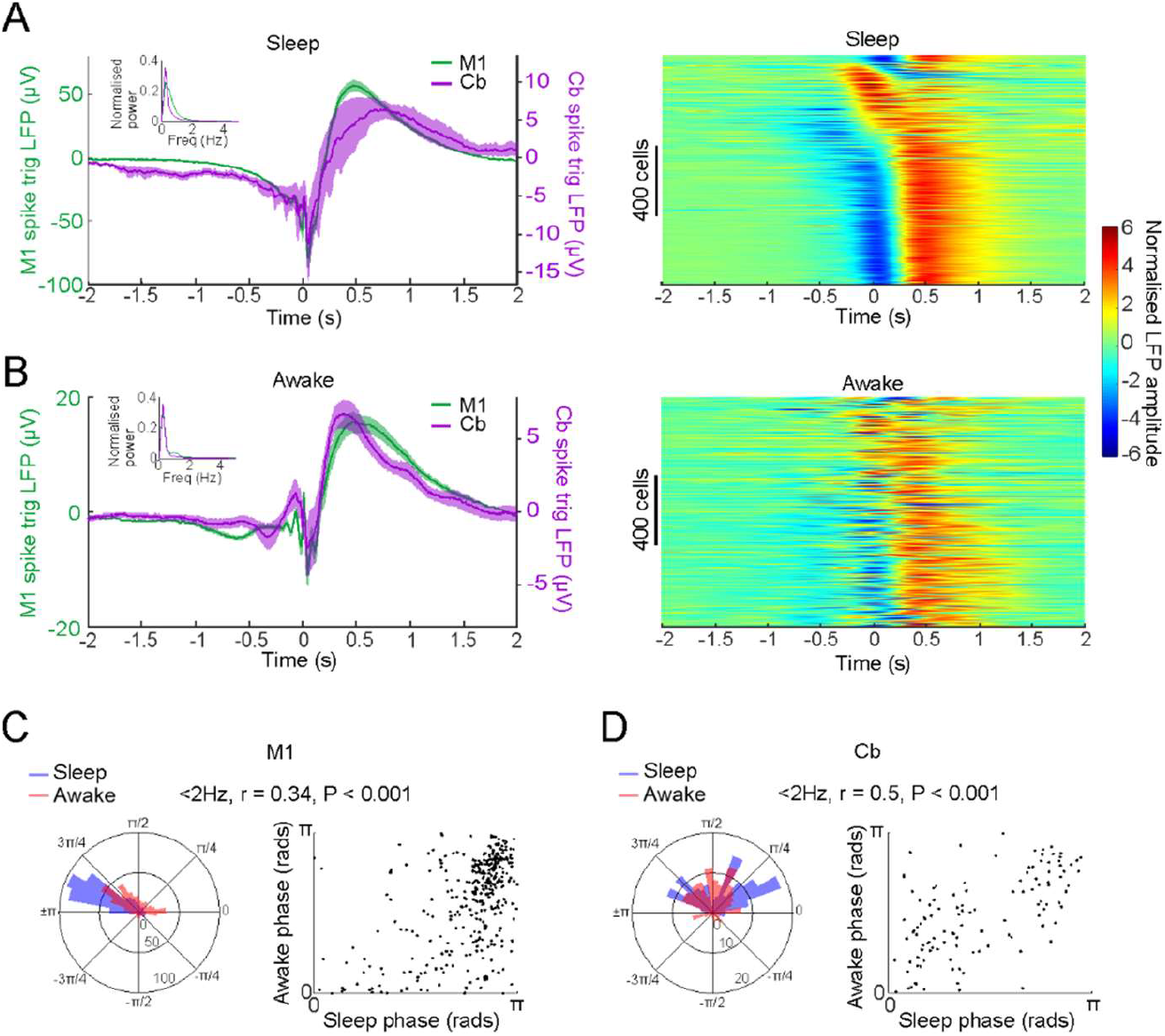
Spike triggered LFP profiles are similar between sleep and awake. A. Left: mean spike triggered M1 LFP during sleep for M1 and cerebellar spikes. Inset shows mean LFP power. Right: all M1 and Cb spike triggered LFPs plotted in ascending order of instantaneous phase at zero lag, normalised by z-scoring. B. Same as A but during awake. Cells are plotted in right panel using the same order as in A. C. Left: polar histogram of all instantaneous phases at t=0s for M1 spike triggered LFPs low pass filtered <2Hz; Right: said phases plotted during sleep against waking with circular-circular correlation coefficient and P value indicated. D. Same as C but for cerebellar spikes.

### Similar low-frequency cell-cell coupling between wake and sleep

Next we examined pair-wise spike cross-correlograms between simultaneously-recorded M1-M1, Cb-Cb and M1-Cb cell pairs (examples shown in Figure 4A-C). Figure 4D-F shows cross-correlograms for all cell pairs, sorted according to the relative peak-time of the low-frequency (<2Hz) correlation structure in sleep. Visual inspection again suggests that, at the population level, there is similarity between the correlation structure in wake and sleep, in that the timing of peaks relative to time-zero (which indicates a tendency for one cell to fire before/after the other) is preserved between brain states. To quantify this, we calculated an index of cross-correlation asymmetry from the difference between the area under the left and right sides of the cross-correlogram (normalised by the total area). We then calculated the linear correlation of this asymmetry measure between wake and sleep across all cell pairs. Figures 4G-I show a significantly positive correlation for M1-M1, Cb-Cb and M1-Cb cell pairs, again demonstrating similarity at the population level between neuronal dynamics in waking and sleep.

**Figure 4.**
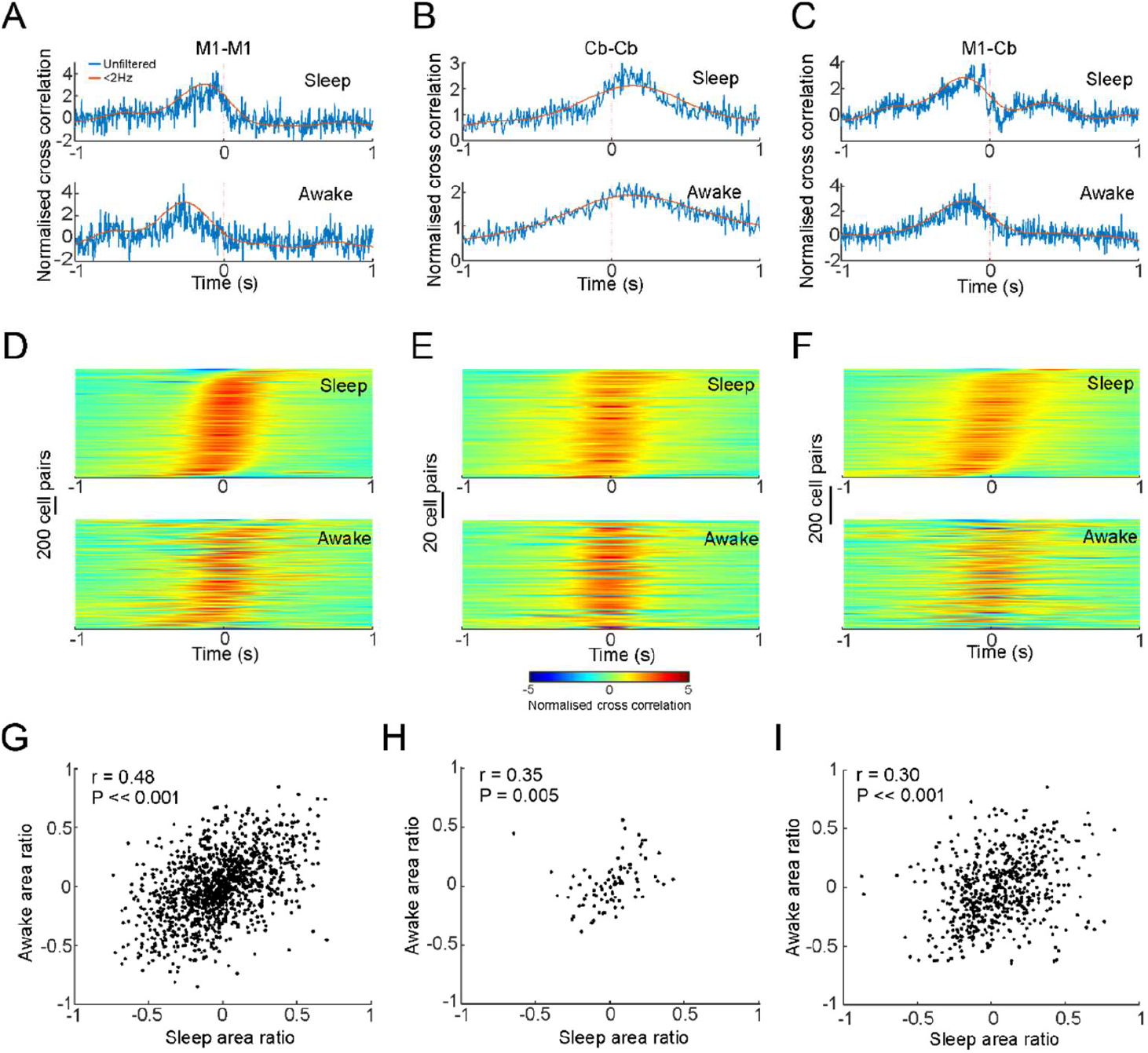
Spike-spike cross correlations in wake and sleep. A-C Example cross-correlations between pairs of M1 and cerebellar neurons during sleep and wake. D. Top: all M1-M1 spiking cross correlations filtered <2Hz and ordered in ascending sequence of relative peak time during sleep. Bottom: waking cross correlations plotted using the same order as sleep. E-F. Same but for Cb-Cb and M1-Cb spike pairs. Cross correlations were normalised by dividing by the number of overlapping samples and then z-scoring. G-I. Correlation between cross-correlogram asymmetry in wake and sleep across all cell pairs. Linear correlation coefficients and P values indicated. See methods for technique of calculating significance.

To gain insight into the frequencies at which the dynamics were similar, we calculated the cross-spectrum and coherence between the spiking activities of simultaneously-recorded cell pairs. The cross-spectrum is the frequency-domain equivalent of the cross-correlogram, and its phase yields information about the relative phase of the activity of two neurons across frequencies. In other words, a positive/negative cross-spectral phase indicates which neuron within a pair is leading/lagging the other at that frequency. Figure 5A plots the cross-spectral phase for an example pair of M1 neurons during waking and sleep. For this example, the relative phase is broadly similar for low frequencies but diverges for higher frequencies.

**Figure 5.**
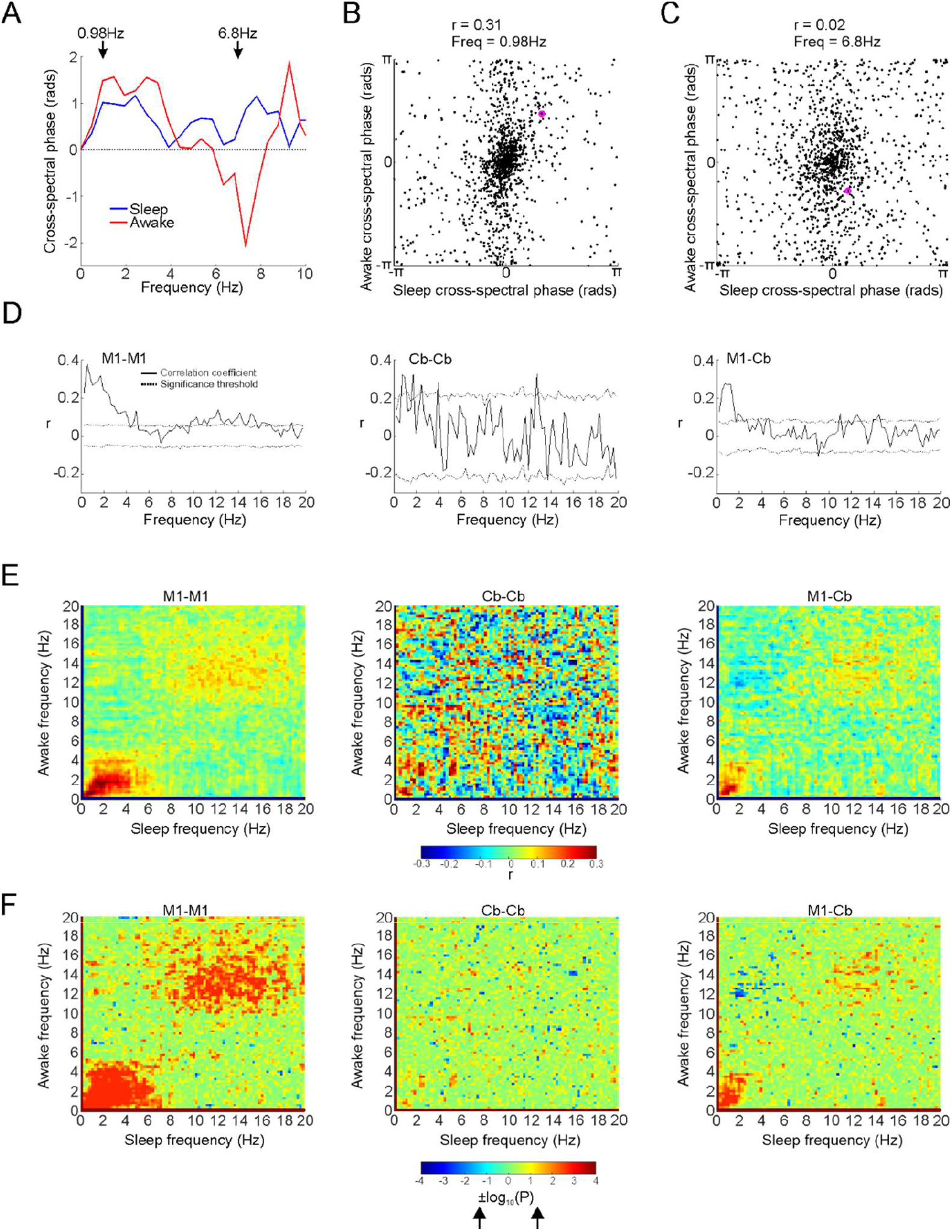
Comparison between sleep and awake spiking phases. A. Cross-spectral phase for an example pair of M1 neurons in sleep and awake. B & C. Sleep vs awake M1-M1 spiking cross-spectral phase at 0.98Hz and 6.8Hz. Example from A circled. D. Circular-circular correlation coefficients between sleep and waking spike cross spectra for each frequency between M1-M1, Cb-Cb and M1-Cb cell pairs. 95% significance thresholds indicated by dotted trace (see methods). E. Circular-circular correlation coefficients between sleep and waking spike cross spectra for all frequency pairs. F. P value for correlation coefficients expressed as ±log10(P). Negative sign gives positive value and correspond to positive r values (and vice versa). Arrow on colour bar indicate values that correspond to P=0.05.

Figures 5B and C show the relative cross-spectral phases during sleep and waking for all M1 pairs at two separate frequencies (0.98Hz and 6.8Hz, indicated by arrows in Fig. 5A). The values corresponding to the example M1 cell pair are circled. Across all pairs, there is a significant correlation between relative phase in waking and sleep at the low frequency, but not at the higher frequency. In other words, at low frequencies, if one cell leads the other during waking activity, it is more likely also to lead during sleep. Figure 5D shows the circular correlation between relative phases of pairwise cross-spectra in waking and sleep for all simultaneously-recorded pairs of M1-M1, Cb-Cb and M1-Cb neurons across all frequencies. We found there was significant similarity in the low-frequency pairwise correlation structure sleep in all cases. Additionally, there was weak but significant positive correlation at spindle-band frequencies for M1-M1 and M1-Cb pairings. To examine whether there were any cross-frequency relationships between waking and sleeping we also performed the circular correlation analysis across different frequencies in waking and sleep (figure 5E,F). However, in all cases, the strongest similarity was seen between low-frequencies in waking and low-frequencies in sleep.

### Correlation structure during up-state events

The above results establish similar low-frequency population structure within and between M1 and Cb in waking and sleep. To further investigate when in sleep this structure occurs, we focussed on identified up- state events in sleep. Figure 6A-C shows spike-spike coherence calculated using a centred sliding window from -3s to 3s around up-state events. Clear increases in coherence at both low-frequencies and spindle frequencies were associated up-state events. Repeating our cross-spectral phase correlation analysis for only time-windows around up-states (Fig. 6D-I) replicated the results of our analysis across all sleep (Figure 5) in revealing a similar correlation structure with waking activity. Therefore, we conclude that up-states are associated with both spindle-frequency coupling between the neocortex and cerebellum, as well as a preserved low-frequency sequential firing structure that is similar to waking activity.

**Figure 6.**
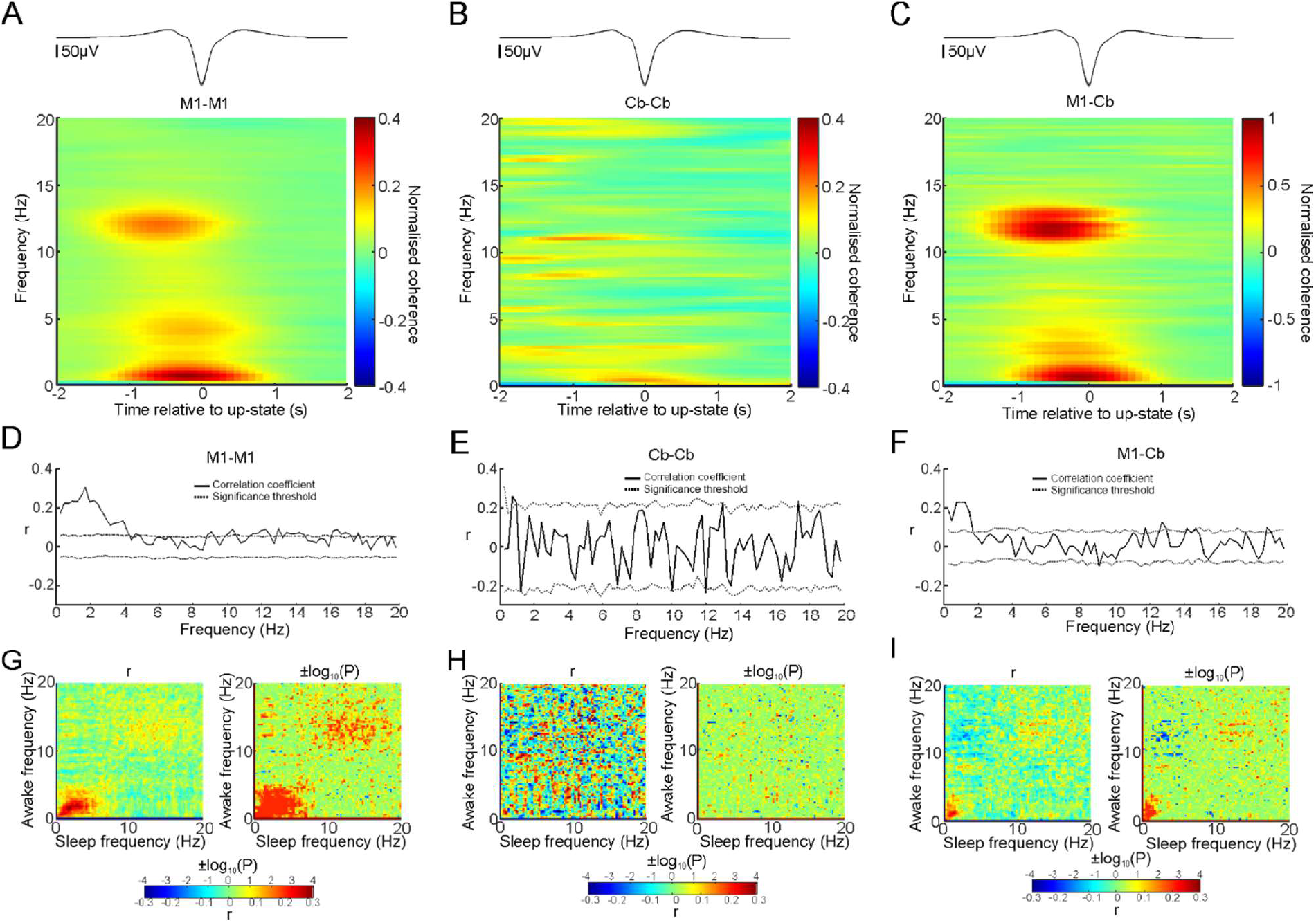
Event aligned sleep and waking spike coherence. A-C Mean M1-M1, Cb-Cb and M1-Cb spike coherence aligned by sleep up-states. Up-state aligned LFP traces are shown above the plot. D-F. Circular-circular correlation coefficients between cell-cell cross-spectral phase calculated using only FFT windows centred over sleep up-states. 95% significance thresholds indicated by dotted trace. G-I. Circular-circular correlation and significance between all frequency pairs for M1-M1, Cb-Cb and M1-Cb cell pair cross-spectra.

### M1 and cerebellar population activity in sleep and awake behaviour

Although the pairwise low-frequency correlation structure between individual neurons is preserved between waking and sleep, our analysis does not exclude the possibility of an overall systematic shift in the timing of population activity in M1 relative to the cerebellum. To address this question explicitly we first examined the grand average cross-correlogram between all pairs of M1-Cb neurons in wake and sleep (Fig. 7A). The peak of the averaged cross-correlation in sleep is for M1 activity leading cerebellar activity, while during waking the correlation appears symmetrical. We quantified this using the average cross-correlation asymmetry across all M1-Cb cell pairs. In sleep, there was a clear and significant asymmetry in the direction of M1 leading the cerebellum (P<0.001, mean asymmetry values calculated for each session, comparison made across sessions). By contrast, in waking periods, there was no significant asymmetry (P = 0.78, Fig. 7B), indicating that there was a similar proportion of cells pairs with M1 leading and lagging the cerebellum.

**Figure 7.**
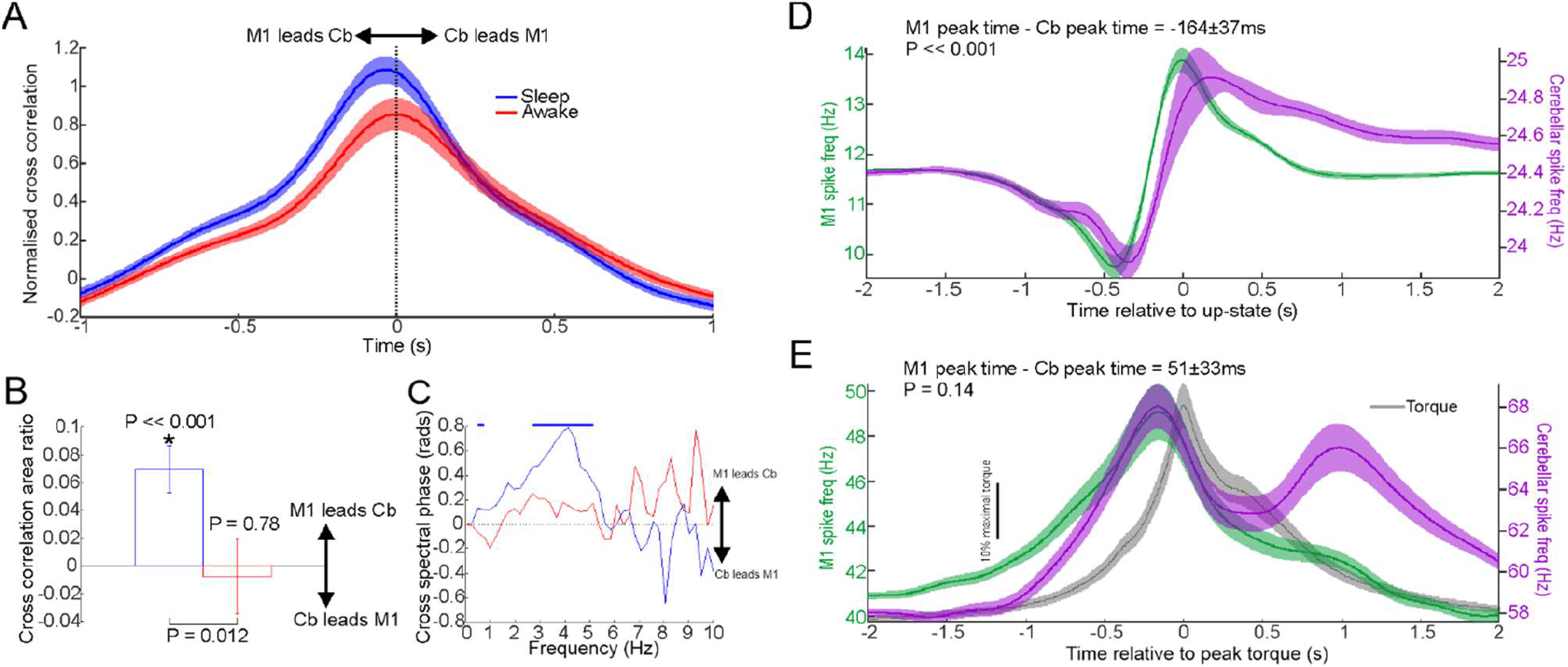
Shift in overall timing of M1 and cerebellar activity between wake and sleep. A. Mean cross-correlation between all pairs of M1 and cerebellar neurons in wake and sleep. B. Average cross-correlogram asymmetry across all pairs in wake and sleep. C. M1-Cb spike-spike cross spectral phase during wake and sleep. Horizontal bars indicate frequencies where phase is significantly different to zero (5% confidence level, circular k-test). D. M1 and Cb spike firing low pass filtered at 2Hz and aligned by sleep up-states. E. Wrist torque, M1 and Cb spike firing low pass filtered at 2Hz aligned by peak wrist torque during waking visuo-motor task.

Next, we examined the grand average cross-spectral phase between all M1-Cb cell pairs. This also confirmed a change in relative phase at low-frequencies between waking and sleep. In sleep, M1 firing led the cerebellum at most frequencies below 5 Hz, while during waking there was no significant mean phase shift between M1 and cerebellar firing (phase for each session derived from averaged cross-spectra within the session, comparison made across sessions, Fig. 7C).

### M1 and cerebellar population activity during up-states and isometric wrist movements

Next, we examined whether the asymmetry between M1 and Cb population activity at low frequencies in sleep was related to the dynamics of down-to-up state transitions. Figure 7D shows average firing rates of all cells in M1 and Cb aligned to up-state events (low pass filtered <2Hz). Although the firing rate profiles are remarkably similar between areas, it is apparent that M1 activity precedes the cerebellum. The time of peak firing differed significantly across the population of cell pairs with M1 leading the cerebellum by 164±37ms (P<0.001, one-sample t-test across sessions).

To obtain a comparable measure of the temporal offset in firing rates associated with movement, we analysed a dataset from monkeys U and Y during performance of an isometric wrist torque task (Hall, de Carvalho et al. 2014). Monkeys were trained to move a cursor from the centre of the screen to one of eight peripheral targets by generating torque at the wrist. We averaged M1 and cerebellar spike firing rate aligned to the time of peak wrist torque (combining all target directions) and low-pass filtered the profiles at 2Hz (Fig.7E). Activity in both M1 and cerebellum consistently preceded the time of peak wrist torque. (Note that cerebellar neurons also exhibited a second later firing peak, possibly related to releasing the torque or receiving food reward.) Across the population, there was no significant difference in the time of the first peak cerebellar activity compared to M1 activity (first peak defined as highest local maxima occurring before 0.5s, P=0.14, one-sample t-test across sessions). Taken together, these results all support a systematic shift in the relative timing of M1 and cerebellar activity from waking (when M1 and cerebellar activity is broadly contemporaneous) to sleep (when M1 firing tends to lead the cerebellum).

## Discussion

Previously we have shown that cerebro-cerebellar functional connectivity in sleep LFPs exhibits a reciprocal pattern of low-frequency causality directed from motor cortex to cerebellum and spindle-frequency causality from the cerebellum, via the thalamus, to M1 (Xu, De Carvalho et al. 2021). This study builds on these findings by examining the structure of population activity in the motor cortex and cerebellum in sleep compared with waking activity. We used time-domain and frequency-domain correlation to show a broad preservation of sequential firing within and across areas between these different brain states. However, we also observed a systematic shift from contemporaneous activity in M1 and cerebellum in waking states to an overall time-lag in sleep, such that up-state events in the motor cortex preceded up-state events in the cerebellum.

Awake activity was characterised by low-frequency, rhythmic peaks in EMG which were correlated with troughs in motor cortex field potentials and elevated firing rates. Previously we have shown that such ‘movement intermittency’ arises from an interaction between extrinsic sensory feedback loops and intrinsic dynamics in motor circuitry, which we suggested may reflect internal models of the body and environment used for predictive control of movement (Susilaradeya, Xu et al. 2019). These intrinsic dynamics are evident from the structure of multiple motor cortical neurons or LFPs which, when projected onto a plane reveal rotational trajectories consistent with a sequential activity pattern (Churchland, Cunningham et al. 2012). Importantly, these rotational patterns are preserved in sleep, particularly around up-state events, suggesting that they arise at least in part from intrinsic connectivity rather than overt waking behaviour (Hall, de Carvalho et al. 2014). Thus, the low-frequency dynamics in up-state activity patterns resemble movement-related activity and may correspond to a kind of ‘fictive’ movement. Consistent with this, we observed that low-frequency sequential firing patterns in motor cortex (characterised by the asymmetry of spike cross-correlograms and the phase firing rate cross-spectra) were broadly conserved between waking and sleep, including around sleep up-states. In addition, using simultaneous recordings in two areas, we were able to show that this was also true for correlations between motor cortex and cerebellum, as well as within the cerebellum itself (despite our smaller sample of simultaneously recorded cerebellar units). Thus, activity in cerebro-cerebellar networks in sleep may be performing similar computations associated with fictive movements even though no overt movement is being generated due to gating at the spinal level.

An influential theory of cerebellar function is that, during waking behaviour, an efference copy of the descending motor command is passed to the cerebellum (possibly via the mossy fibre system) and is used to predict the consequences of movement via internal models. These predictions then drive corrective movements via output connections back to motor cortex via the thalamus. Such reciprocal connectivity may explain why activity in M1 and the cerebellum is broadly contemporaneous in our visuomotor task and during free behaviour. By contrast, in sleep, cerebellar activity was delayed relative to motor cortex, evidenced by asymmetrical cross-correlograms and non-zero cross-spectral phase at low frequencies. The delay between motor cortical and cerebellar activity was most apparent in the average firing rate for each area relative to sleep up-states, suggesting up-state events are initiated in the neocortex and then propagate to the cerebellum, consistent with previous observations (Rowland, Goldberg et al. 2010, Xu, De Carvalho et al. 2021). This asymmetry, as well as the reduced low-frequency coherence between M1 and cerebellum in sleep, may be explained by the altered state of the thalamus reducing the low-frequency connectivity from the cerebellum to neocortex in sleep. Thus, at low frequencies in sleep, the causal influence is predominantly from neocortex to cerebellum. However, up-state events are also associated with increased cerebro-cerebellar spike-spike coherence at spindle frequencies, and our previous study showed that the direction of causal influence at these frequencies was from the cerebellum to neocortex (Xu, De Carvalho et al. 2021). Thus, it seems that although connectivity between the cerebellum and neocortex is altered, spindle oscillations provides a mechanism by which the cerebellum can nevertheless influence motor cortex in sleep. Notably, this is a frequency band that is known to be effective at driving neocortical plasticity (Rosanova and Ulrich 2005).

These observations lead us to propose the following speculative hypothesis. During waking movements, the cerebellum generates predictions of the consequences of actions which, combined with actual sensory inputs, guide corrective movements. This reciprocal connectivity leads to strong low-frequency coherence with approximately zero phase-lag. Motor cortex activity during up-state events reflects fictive movements, and the cerebellum similarly generates predictions of the fictive consequences of these actions. However, these predictions do not generate corrective movements, leading to unidirectional cortico-cerebellar connectivity and delayed cerebellar activity. However, cerebellar activity is relayed at spindle-frequencies back to the motor cortex, and may thus provide a training signal to guide plasticity in motor cortex. Such a mechanism would provide a means for short-term learning stored in internal models in the cerebellum to be consolidated in long-term changes in motor cortical networks. Moreover, such a mechanism of fictive practice during sleep, guided by predicted consequences computed by the cerebellum, could explain how performance can even improve during sleep (Schonauer, Geisler et al. 2014). These speculations require further experiments to test. In particular, it will be interesting in future to examine how activity patterns in motor cortex and cerebellum in sleep are influenced by previous waking experience. Such experiments will hopefully shed more light on the mechanisms through which ‘practice with sleep makes perfect’.

## Acknowledgments

This work was supported by the Wellcome Trust (106149) and the Engineering and Physical Sciences Research Council (H051570).

The authors would like to thank Mr Norman Charlton and Ms Jennifer Tulip for their technical assistance.

